# A metal artifact reduction method for small field of view CT imaging

**DOI:** 10.1101/2019.12.27.889204

**Authors:** Seungwon Choi, Seunghyuk Moon, Jongduk Baek

## Abstract

To reduce metal artifacts, several sinogram inpainting-based metal artifact reduction (MAR) methods have been proposed where projection data within the metal trace region of the sinogram are treated as missing and subsequently estimated. However, these methods generally assume data truncation does not occur and all metal objects reside inside the field-of-view (FOV). For small FOV imaging, these assumptions are violated, and thus existing inpainting-based MAR methods would not be effective. In this paper, we propose a new MAR method to reduce metal artifacts effectively in the presence of data truncation. The main idea behind the proposed method is the synthesis of a sinogram, which is treated as the originally measured sinogram. First, an initial reconstruction step is performed to remove truncation artifacts. The next step consists of a forward projection of the small FOV image, replacing the original sinogram with the synthesized sinogram. The final step is the application of sinogram inpainting based MAR methods using the synthesized sinogram. Verification of the proposed method was performed for three situations: extended cardiac-torso (XCAT) simulation, clinical data, and experimental data. The proposed method was applied with linear MAR (LMAR) and normalized MAR (NMAR), and the performance of the proposed method was compared with that of the previous method. For quantitative evaluation, normalized mean squared error (NMSE) and structure similarity (SSIM) were used. The results show the effectiveness of the proposed method to reduce residual metal artifacts, which are present in the results obtained with the previous method. The evaluation results using NMSE and SSIM also indicate that the proposed method is more effective than the previous method.

## Introduction

During X-ray computed tomography (CT) imaging, the presence of metallic implants such as dental fillings, hip prosthesis, and orthopedic implants introduces metal artifacts in the reconstructed images. The high atomic number of metals leads to photon starvation, beam hardening, and scatter, which cause severe dark and bright streak artifacts in the reconstructed CT images [1, 2]. These metal artifacts degrade the image quality severely and interrupt the diagnostic performance.

To reduce the metal artifacts, several metal artifact reduction (MAR) algorithms have been proposed. One of the most widely known methods, called sinogram inpainting, finds metal trace region within a sinogram corrupted by metal objects and replaces them with the appropriate values. There are various methods to fill in metal trace region such as simple linear interpolation using peripheral values known as linear MAR (LMAR) [3], high-order [4–6], wavelet [7, 8], and prior knowledge [9, 10]. However, these methods are effective only for simple structures and may introduce additional artifacts due to the increase of estimation errors when applied to images containing complex anatomical structures. To reduce the estimation errors, normalized MAR (NMAR) [11] was proposed, where prior images were used to minimize the estimation errors in the metal trace region. However, the NMAR method estimates the prior image from an LMAR image. Since the accuracy of the prior image varies depending on the quality of the LMAR image [11], when severe metal artifacts are present, distortion may occur in the prior image itself, and thus additional artifacts would appear in the corrected image.

With a general sinogram inpainting method, metal is segmented from an uncorrected image and then forward projected to identify the metal trace region in sinogram. While inpainting-based methods are effective in MAR, they assume object truncation does not occur and metal objects reside inside the field-of-view (FOV). However, when the FOV is small and multiple metals are present outside the FOV such as dental CT imaging, or when the metal object is not covered by the FOV such as bone biopsy needles, the basic assumption of the existing inpainting-based MAR algorithms is violated. If metal objects reside outside the FOV, the reconstructed image within the FOV can not be used to identify the location of metal outside the FOV, and thus the metal trace region can not be found completely in the sinogram. In addition, for small FOV imaging, projection data contain information about the materials present outside the FOV. With the sinogram inpainting method using a prior image, the value of the prior sinogram is mismatched with the value of the uncorrected sinogram and interferes with the MAR algorithms because the prior image only represents the small FOV. Thus, for metals that reside outside the FOV, the existing sinogram inpainting based MAR methods would not be effective.

In this work, we propose a new method to reduce metal artifacts for small FOV imaging in the presence of data truncation. The proposed method conducts MAR by using a sinogram of the FOV obtained through the forward-projection of an uncorrected metal artifact image that has undergone only truncation correction. The proposed method was validated using simulated extended cardiac-torso (XCAT) data, clinical data, and an experimental cylinder phantom data containing metal implants outside the FOV. The proposed method was applied to LMAR and NMAR as a sinogram inpainting method, and its performance was compared with that of the previous method. For quantitative evaluation, the normalized mean squared error (NMSE) and structure similarity (SSIM) were used.

## Methods

### Proposed Method

The basic idea of the proposed method is to synthesize the projection data for a small FOV object by conducting forward-projection of the initially reconstructed FOV image, then treating the synthesized data as the originally measured projection data. Although the metal artifacts appear throughout the entire image, artifacts are confined within the metal trace region of the sinogram after the forward projection. The overall diagram of the proposed method is depicted in Fig. 1 and the detail of each step is as follows.

**Fig 1.**
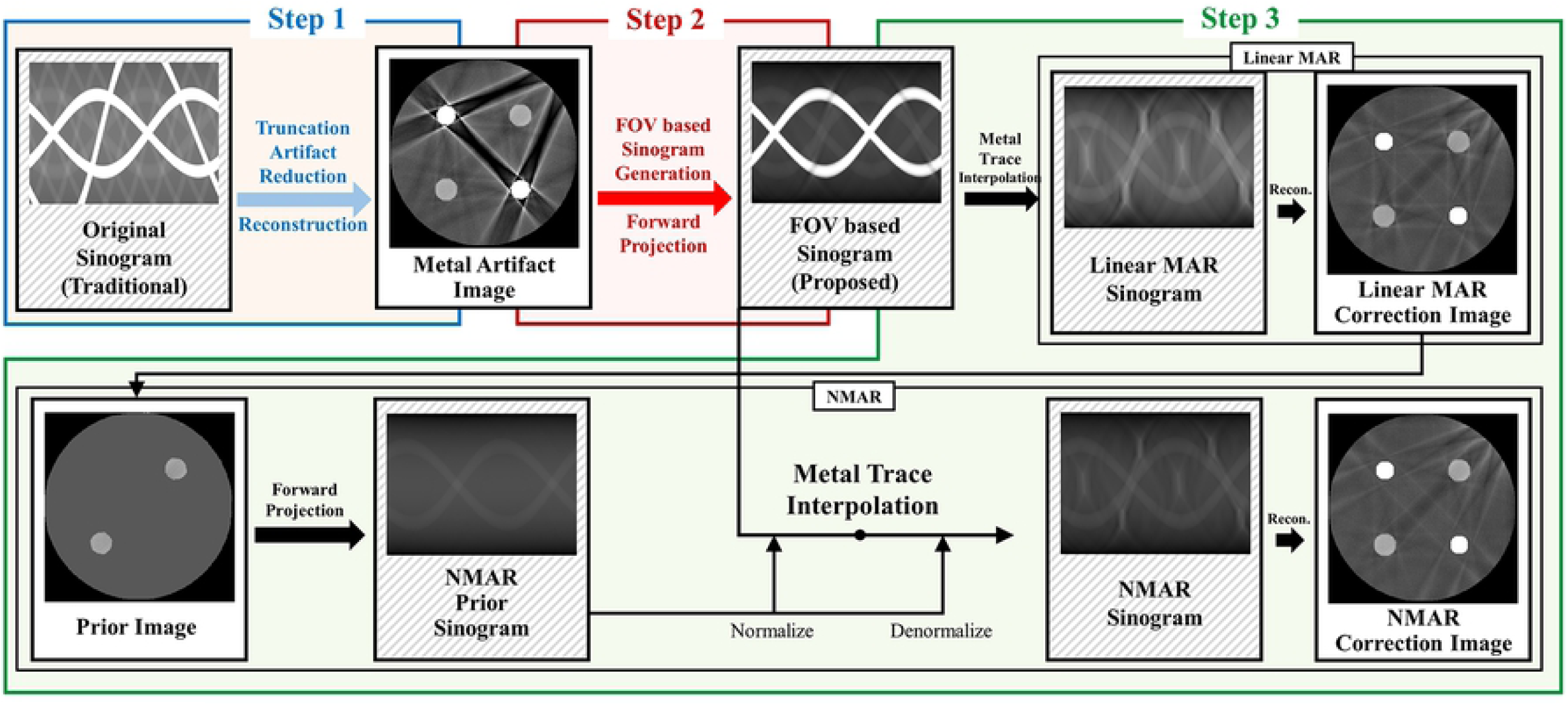
Diagram of the proposed method.

The first step is to conduct a small FOV reconstruction (Step 1). The original uncorrected sinogram has missing information because of the data truncation.

Truncation artifacts occur when the sinogram is reconstructed without any processing, so truncation correction is necessary to reduce the truncation artifacts in small FOV image. To compensate for the truncation artifacts, we use the symmetric mirroring method [12] to estimate the projection data near the truncated region in the sinogram space. The symmetric mirroring method estimates the truncated sinogram region by deducing the data inside and outside the FOV as symmetric. In order to estimate projection data within the truncated region, we symmetrically flip the inner sinogram and multiply the weighting function, which is smoothly reduced to zero. In this work, we use a cosine function as a weighting function. After the first truncation correction, no more truncation artifacts occur in the subsequent process, since the proposed algorithm conducts the forward projection over a small FOV image.

The next step is to conduct a forward-projection of the small FOV image, replacing the original sinogram with the synthesized sinogram (Step 2). Note that forward-projection in step 2 was conducted using five times finely sampled detector size compared to the original detector size in order to reduce the discretization errors in the forward-projection step.

The final step is to apply a sinogram inpainting-based MAR method (i.e. LMAR and NMAR) using the synthesized sinogram (Step 3). For LMAR and NMAR, we use a simple thresholding technique to segment the metal image in the reconstructed small FOV image, and then find the metal trace region in the sinogram space. A prior image of NMAR is created by simple thresholding of the LMAR results. For NMAR, the bone pixel is used as is, and the soft tissue is filled with a single value. All interpolations used in this paper are linear interpolations in the radial direction.

### XCAT Data

To validate the proposed algorithm, the head, shoulder, and hip were simulated with the XCAT phantom image developed by Segars [13]. In order to acquire a metal-inserted image, the metal was created as a binary image by setting the position and shape similar to the metal in a real human body, and the metal was inserted into the original XCAT image. Fig. 2 shows the phantom and the first reconstruction result. In each column, original XCAT images, metal implant binary images, initial reconstruction images, and images with truncation artifact correction are shown.

**Fig 2.**
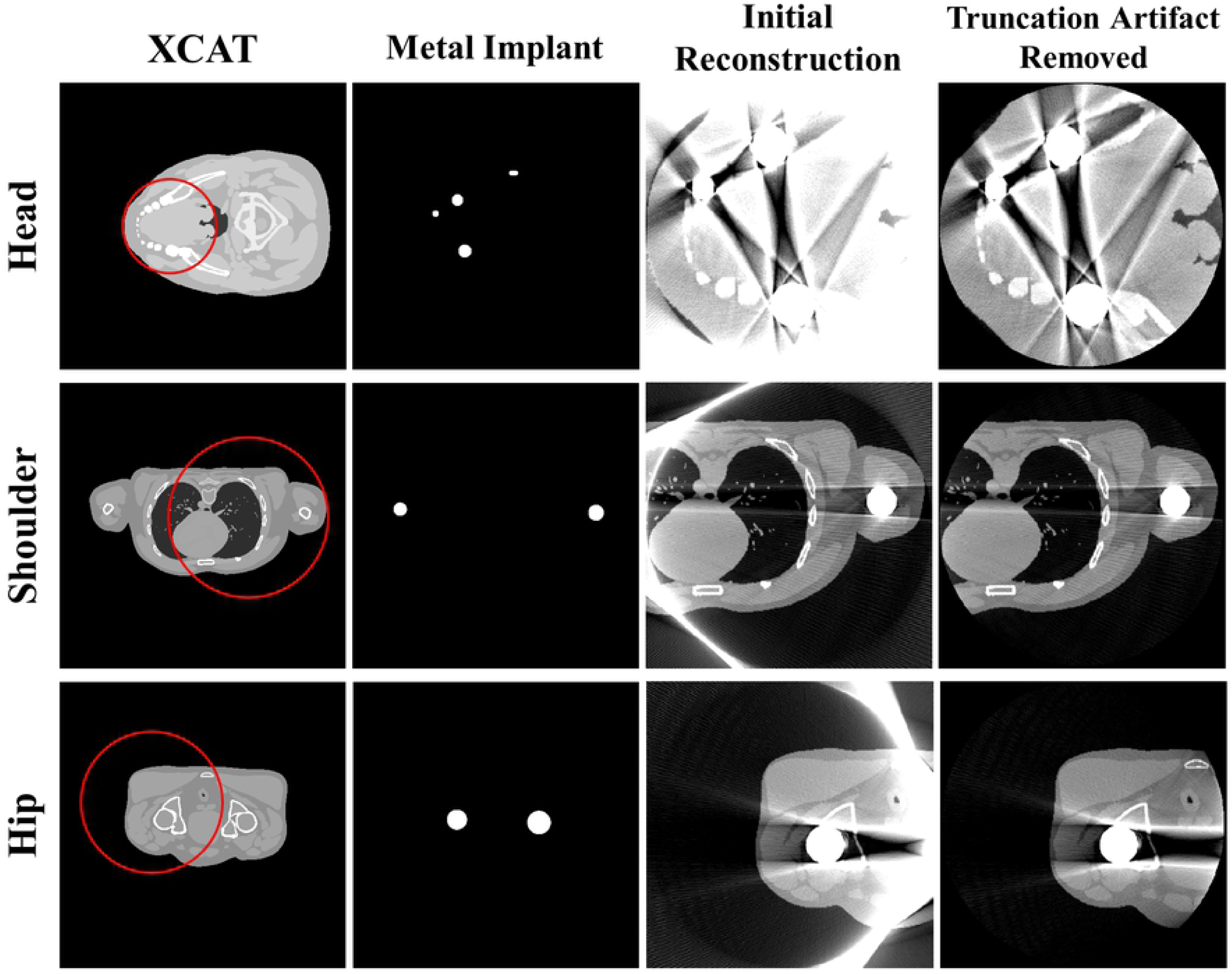
Representative images of XCAT data. Each row corresponds to different body section. Each column represents the original XCAT images, metal implants, reconstruction images, and reconstruction images with truncation artifact correction. In the XCAT images, the red circles represent the FOV for each case. Window width/window level = 2000HU/0HU.

In the simulation, the projection data were acquired using a polychromatic energy spectrum as

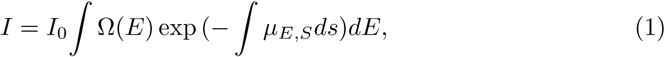

where *I* represents the transmitted intensity and *I*_0_ represents the incident intensity of polychromatic energy. *µ*_*E,S*_ represents the linear attenuation coefficient of the object for each energy and *Ω* (*E*) represents the incident X-ray spectrum for each energy. Polychromatic projection data were acquired with a 120 kVp tube voltage using the Siemens X-ray spectrum [15]. A detailed plot of the X-ray spectrum is shown in Fig. 3. The projection was calculated as

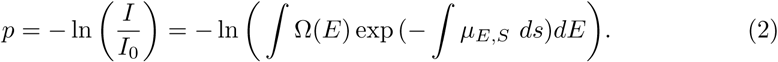

**Fig 3.**
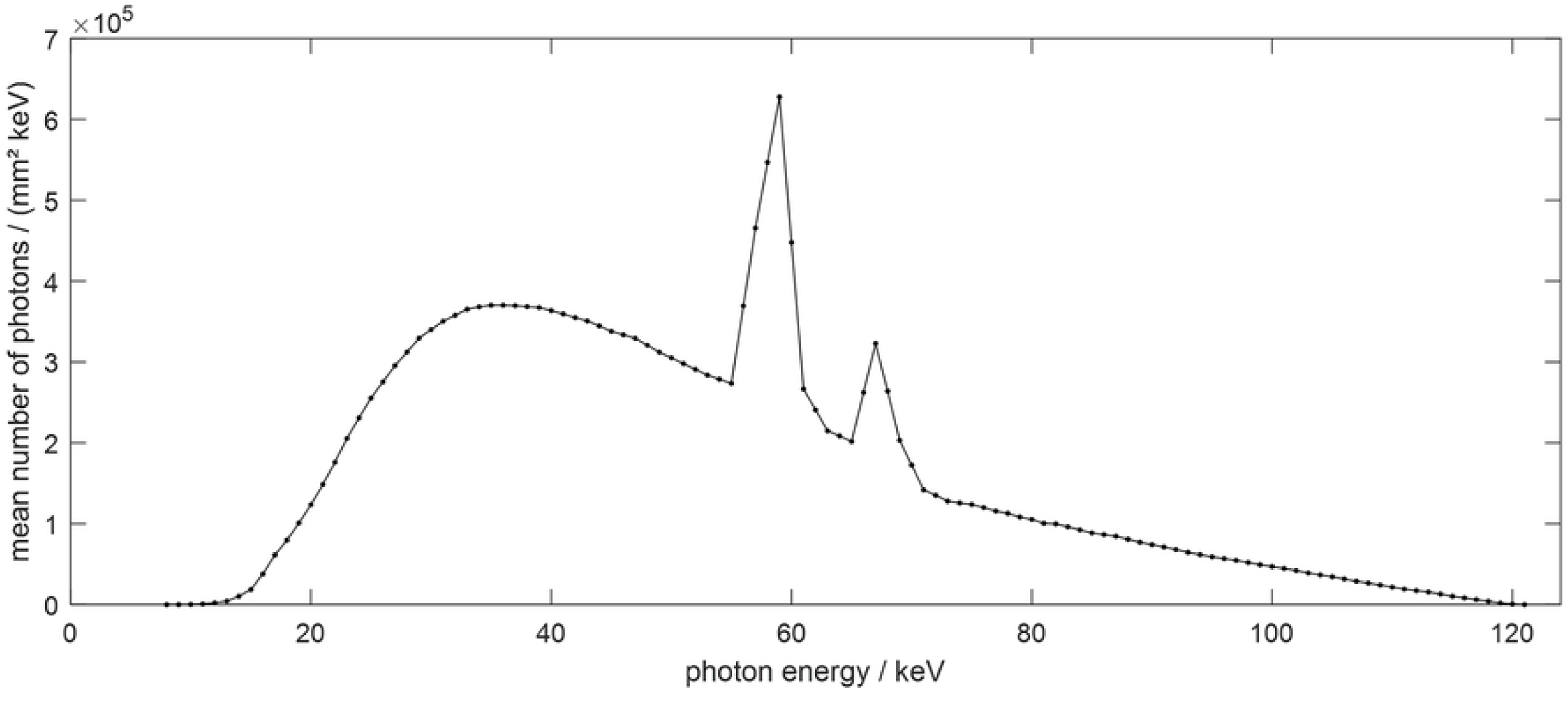
Siemens X-ray spectrum at 120kVp.

In the XCAT simulation, water, bone, and metal were the main components of the object, and thus Eq. (2) can be expressed as

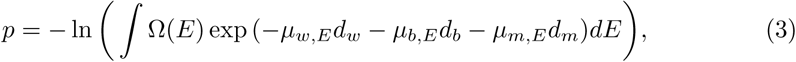

where, *µ*_*w,E*_, *µ*_*b,E*_, and *µ*_*m,E*_ are the attenuation coefficients of water, bone, and metal at a specific energy band *E*, which were obtained from the NIST X-ray attenuation database [16]. *d*_*w*_, *d*_*b*_, and *d*_*m*_ are the path-length of the water, bone, and metal.

In order to simulate the polychromatic projection, it is necessary to know the composition of materials in each pixel. Therefore, the XCAT phantom image should be divided into individual images according to the material type. First, the inserted metal was excluded by simple thresholding and the XCAT data were divided into bone and soft tissue images using pixel values from the remaining images. Since each pixel of the XCAT data did not contain a mixture of different tissues, simple thresholding was sufficient to segment metal, soft tissue, and bone images. Then, using the material-specific images, the corresponding attenuation value and the polychromatic spectrum were applied to Eq. (3) to obtain polychromatic projections. Poisson noise was added to the obtained polychromatic projections.

Forward-projection and reconstruction were conducted using the TIGRE: Matlab-GPU Toolbox [17]. The detailed parameters for XCAT and subsequent simulations are summarized in Table 1. Two different geometries were used to confirm the algorithm works for both conventional and dental CT systems. The reconstructed image size was 512 × 512 and the truncation occurred with the corresponding geometries. The diameter for the FOV was the same as the size of the reconstructed image shown in the Table 1. Each detector cell received **1**0^5^ photons and Poisson noise was added. In order to verify the effectiveness of the algorithm, the initial object was cropped to fit the FOV size to prevent truncation errors. Then, the reference image was reconstructed from the generated polychromtic projection data.

**Table 1.** Parameters for data acquisition and reconstruction in both simulation and experiment

| Parameters | Fab-Beam CT Mode | Dental CT Mode |
| --- | --- | --- |
| Source to iso-center distance | 700 mm | 500 mm |
| Detector to iso-center distance | 500 mm | 200 mm |
| Number of views | 720 view | 720 view |
| Detector cell size | $0.388 \times 0.388 \text{ mm}^2$ | $0.388 \times 0.388 \text{ mm}^2$ |
| Detector number | 1024 | 512 |
| Reconstructed image size | $358.4 \times 358.4 \text{ mm}^2$ | $143.36 \times 143.36 \text{ mm}^2$ |
| Reconstructed matrix size | $512 \times 512$ | $512 \times 512$ |
| Reconstructed pixel size | $0.7 \times 0.7 \text{ mm}^2$ | $0.28 \times 0.28 \text{ mm}^2$ |

### Clinical Data

Simulation using clinical data was conducted to evaluate the performance of the proposed method. We used reconstructed image data from “The 2016 Low-dose CT Grand Challenge” dataset [18] and the cancer imaging archive [19]. Abdomen, shoulder, hip, and head images were selected as shown in Fig. 4. Abdomen, shoulder, and hip cases had geometry similar to the general fan-beam CT geometry, and the head case had geometry similar to dental CT geometry as described in Table 1. In order to generate metal artifacts, metal implants were inserted into the clinical reconstruction images. By following the same procedure to generate polychromatic projection data in XCAT simulation, we created a binary image of the metal implants that was similar to the metals existing in a real human body and replaced the corresponding part in the clinical data. The metal-inserted image was reconstructed using generated polychromatic projection data, and then the proposed algorithm was applied to reduce the metal artifacts.

**Fig 4.**
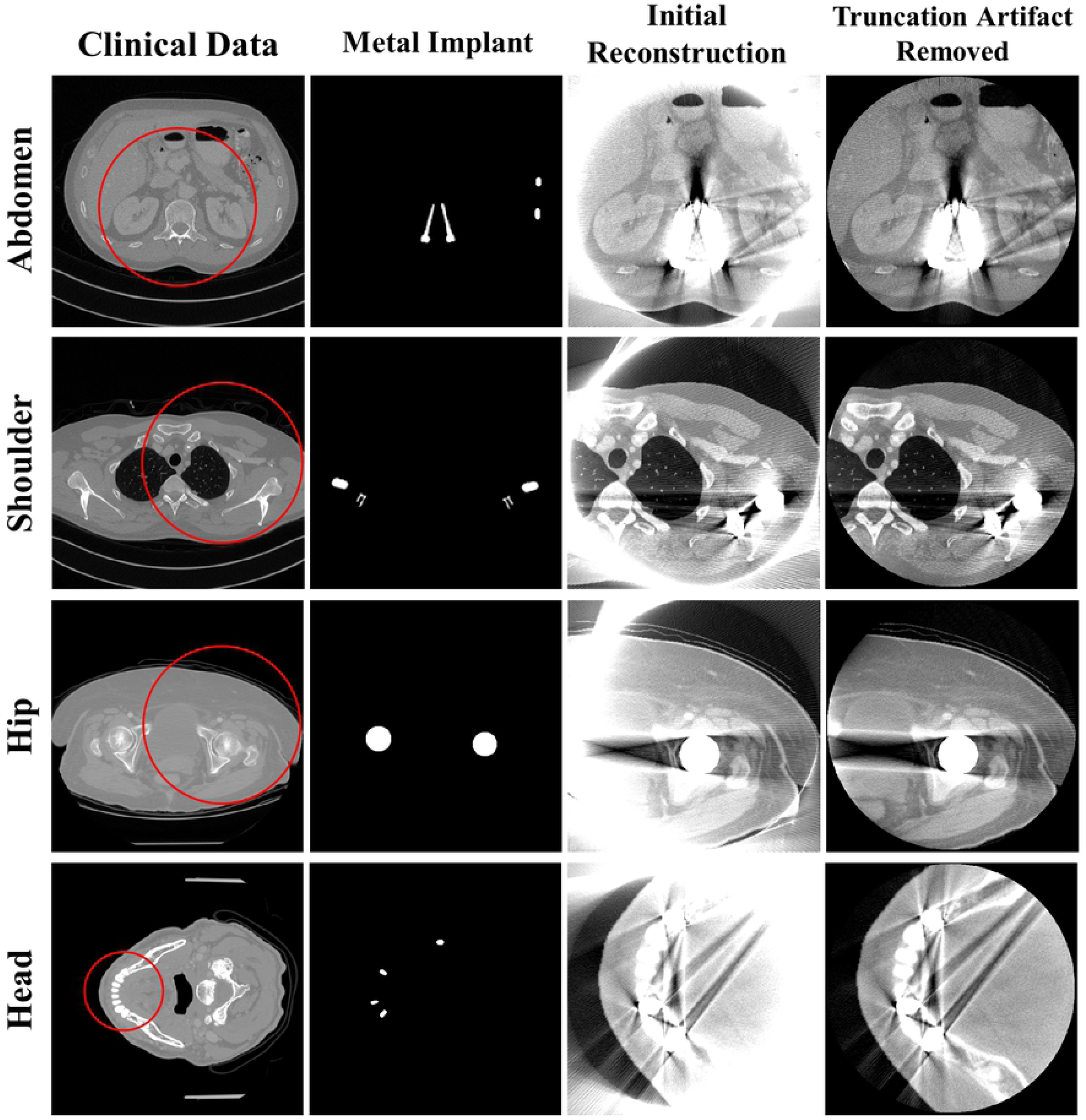
Representative images of clinical data. Each row corresponds to different body section. Each column represents the original clinical images, metal implants, reconstruction images, and reconstruction images with truncation artifact correction. In the clinical data, the red circles represent the FOV for each case. Window width/window level = 2000HU/0HU.

Unlike XCAT data, clinical image is not classified by pixel values precisely for each composing material. Therefore, in order to generate polychromatic projection, the pixel values were given as a mixture of materials by considering the ratio of each component. First, in the images for all materials except for metal, if the pixel value *x* ^*i*^ was less than the threshold *T*_1_(i.e., 80HU), the area was considered soft tissue, and if it was larger than the threshold *T*_2_(i.e., 660HU), the area was considered to be bone. If the value of the pixel was between *T*_1_ and *T*_2_, the area was estimated a mixture of water and bone, and the ratio was obtained using a soft threshold-based weighting method as in Eq.(4) [14]. Bone and water-equivalent tissue components are written as *x*_*b*_ and *x*_*w*_, and the weighting function for each pixel is expressed as follows.

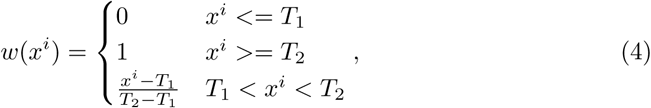

where *w* (*x*^*i*^) is the *i*^*th*^ pixel weighting value. Hence, *i*^*th*^ pixel components of bone and water, 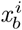 and 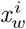, are expressed as

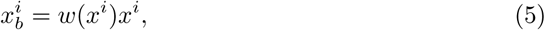

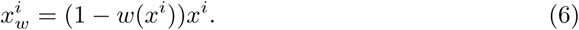

For the bone, water, and metal images, the attenuation coefficient for each energy band was applied according to the spectrum at 120 kVp. We used the same method to generate the polychromatic projection described in previous XCAT data section. To compare the performance of the algorithm in clinical data, a reference image was created by following the same procedure for generating reference XCAT data.

### Experiments

For the experimental data acquisition, we used a benchtop system as shown in Fig. 5(a). The geometry parameters for the benchtop system were identical to the fan-beam CT geometry in the XCAT and clinical data, as shown in Table 1. We conducted the experiments using the disk phantom shown in Fig. 5(b). The disk phantom was 25cm in diameter, and 32 cylinders with 5mm diameters were inserted along its perimeter. In order to examine the robustness of the proposed method, we experimented three cases where the number of metals outside the FOV increase from one to three. Note that the disk phantom contains three metals inside the FOV.

**Fig 5.**
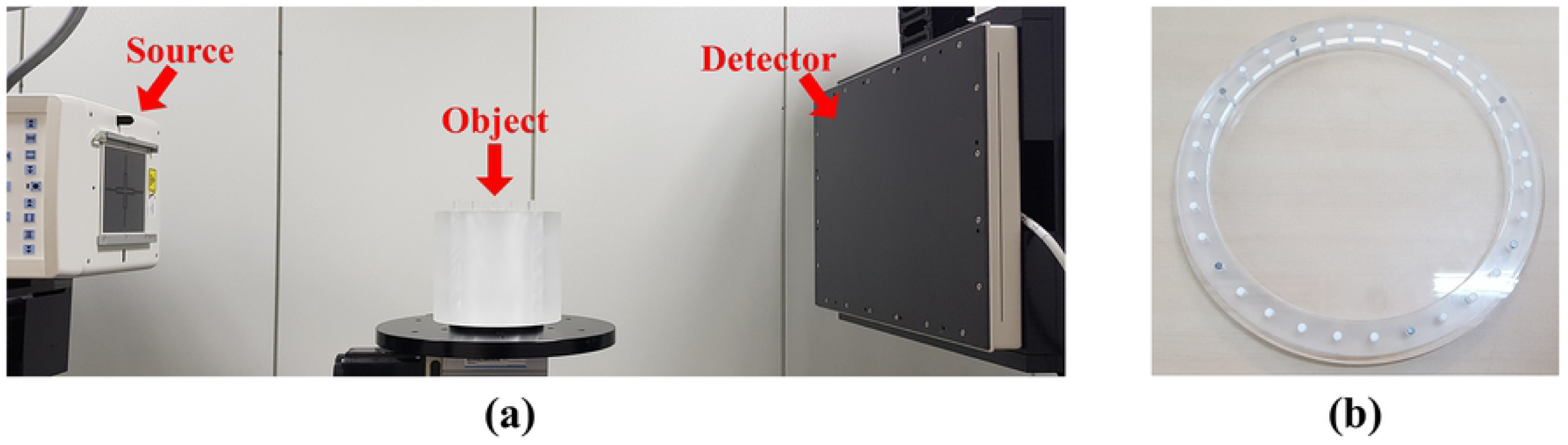
(a): Experimental bench-top system and (b): semi-top view of disk phantom.

We obtained the projection data using a benchtop CT system, which consisted of a generator (Indico 100, CPI Communication and Medical Products Division, Georgetown Ontario, Canada), a tungsten target X-ray source (Varian G-1592, Varian X-ray Product, Salt Lake City, UT) with a 0.6 × 0.6 mm^2^ focal spot and a 300 × 400 mm^2^ flat-panel detector (PaxScan 4030CB, Varian Medical Systems, Salt Lake City, UT). The projection data for the disk phantom were obtained at 90kVp and 6mA. The FDK reconstruction was performed on 512 × 512 matrices with a pixel size of 0.7 × 0.7 mm^2^.

### Image Quality Evaluation

To compare the image quality of the previous and proposed methods with LMAR and NMAR, NMSE and SSIM [20] were used for quantitative evaluation. NMSE is defined as

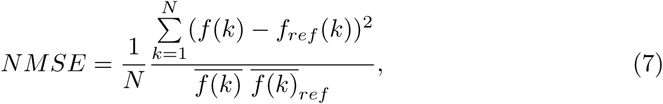

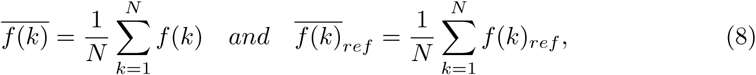

where *f* (*k*) and *f*_*ref*_ (*k*) represent the intensity of the reconstructed and reference images at pixel location *k*, respectively. *N* represents the number of image pixels, and 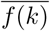 and 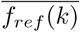 represent the average intensity of the reconstructed and reference imagee, respectively.

SSIM is defined as

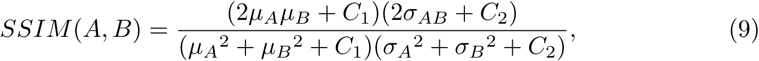

where *µ*_*A*_, *σ*_*A*_, *µ*_*B*_, and *σ*_*B*_ represent the average intensity value and standard deviation of reconstructed images *A* and *B*, respectively. *σ*_*AB*_ represents the covariance between images *A* and *B*, and *C*_1_ and *C*_2_ denote the coefficients for the SSIM calculation ranging from 0 to 1. The NMSE and SSIM were calculated between the reference image and corrected MAR images.

## Results

### XCAT Data Results

Fig. 6 shows the results of previous and proposed LMAR and NMAR methods using XCAT data. Each row corresponds to the head, shoulder, and hip from the XCAT data.

**Fig 6.**
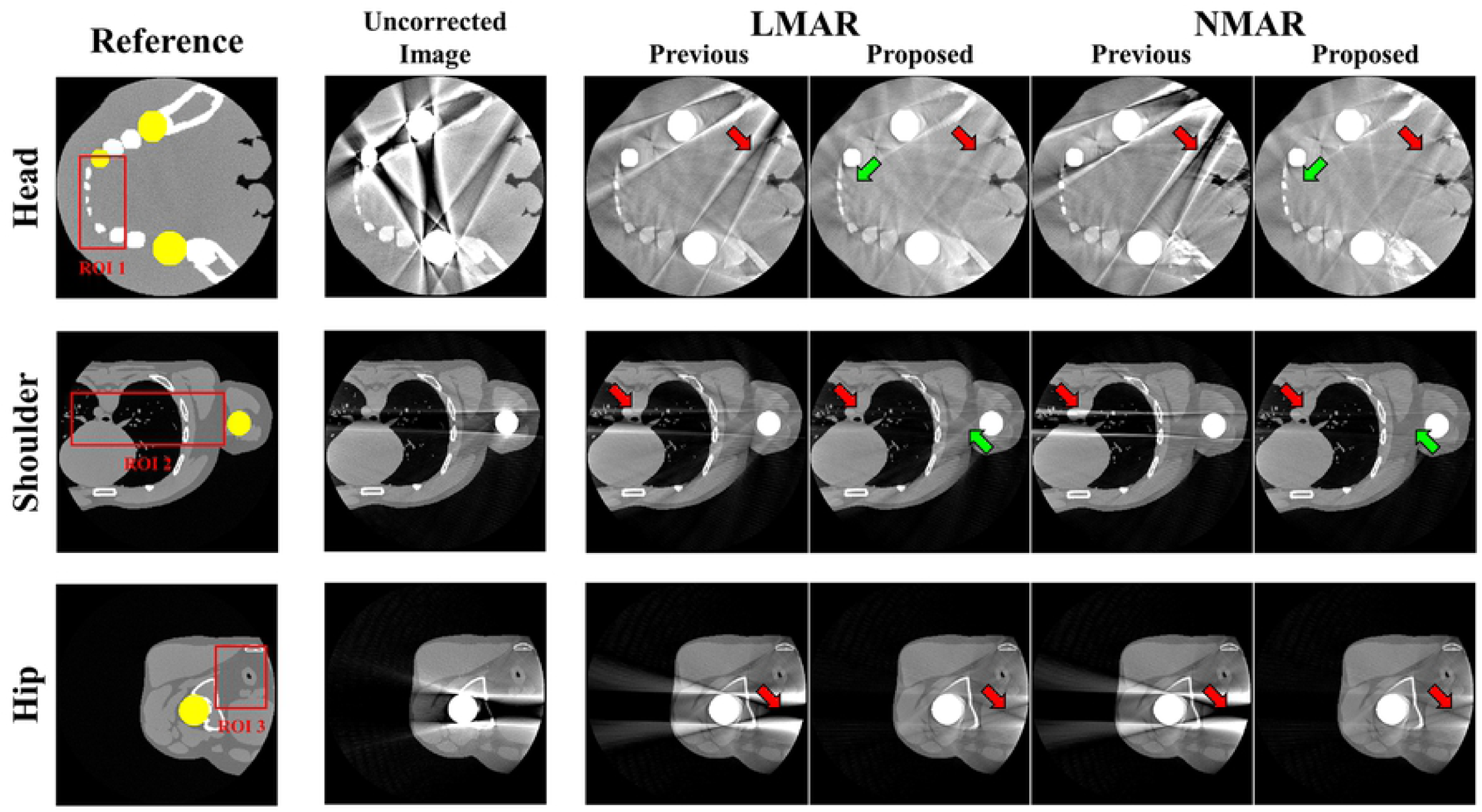
Result with XCAT data. Each row corresponds to different body sections. Each column represents reference images, uncorrected metal artifact images, and previous and proposed LMAR and NMAR result images. Window width/window level = 2000HU/0HU.

The leftmost column shows the reference image for each case, which is an image taken by creating an object without truncation and metals. In each reference image, the yellow area represents the metal object region, and the red box represents the area where the NMSE and SSIM are measured for image quality evaluation. The resulting images show that all MAR processes can reduce metal artifacts compared to the original uncorrected image. However, as indicated by the red arrows, the proposed method provides superior performance for both LMAR and NMAR compared to the previous methods. The metal artifacts produced by the metals inside the FOV can be effectively reduced by both the previous and proposed methods as shown in head image. However, in cases of metal artifacts produced by external metals, previous LMAR and NMAR methods are not effective in removing the artifacts. However, with the proposed method, both LMAR and NMAR could reduces metal artifacts effectively. It is also shown that the NMAR reduces metal artifacts more effectively than LMAR as indicated by the green arrows in Fig. 6.

Fig. 7 shows the sinogram used with the previous and proposed methods for the XCAT-shoulder images and the projection data in the 189th view. In addition, the resulting sinogram from the previous and proposed LMAR and NMAR methods are shown. In the central profile graph, the blue line is the value of each result sinogram and the red line is the value of the reference sinogram. The gray shaded region of the plot shows the metal trace region where interpolation is conducted. It is clear that the proposed method estimates the reference sinogram better than the previous method. The proposed method starts by using the sinogram created through forward-projection of the FOV in Fig. 7(b) instead of the original sinogram. Thus, the value outside the metal trace region is matched well with the reference sinogram. In particular, it is observed that the sinogram with previous method is corrupted by the external metal trace during interpolation, which is prevented effectively by the proposed method. In addition, the previous NMAR method amplifies the estimation error within the metal trace region when the area corrupted by the metal outside the FOV is normalized and denormalized, which adversely affects performance worse than previous LMAR method.

**Fig 7.**
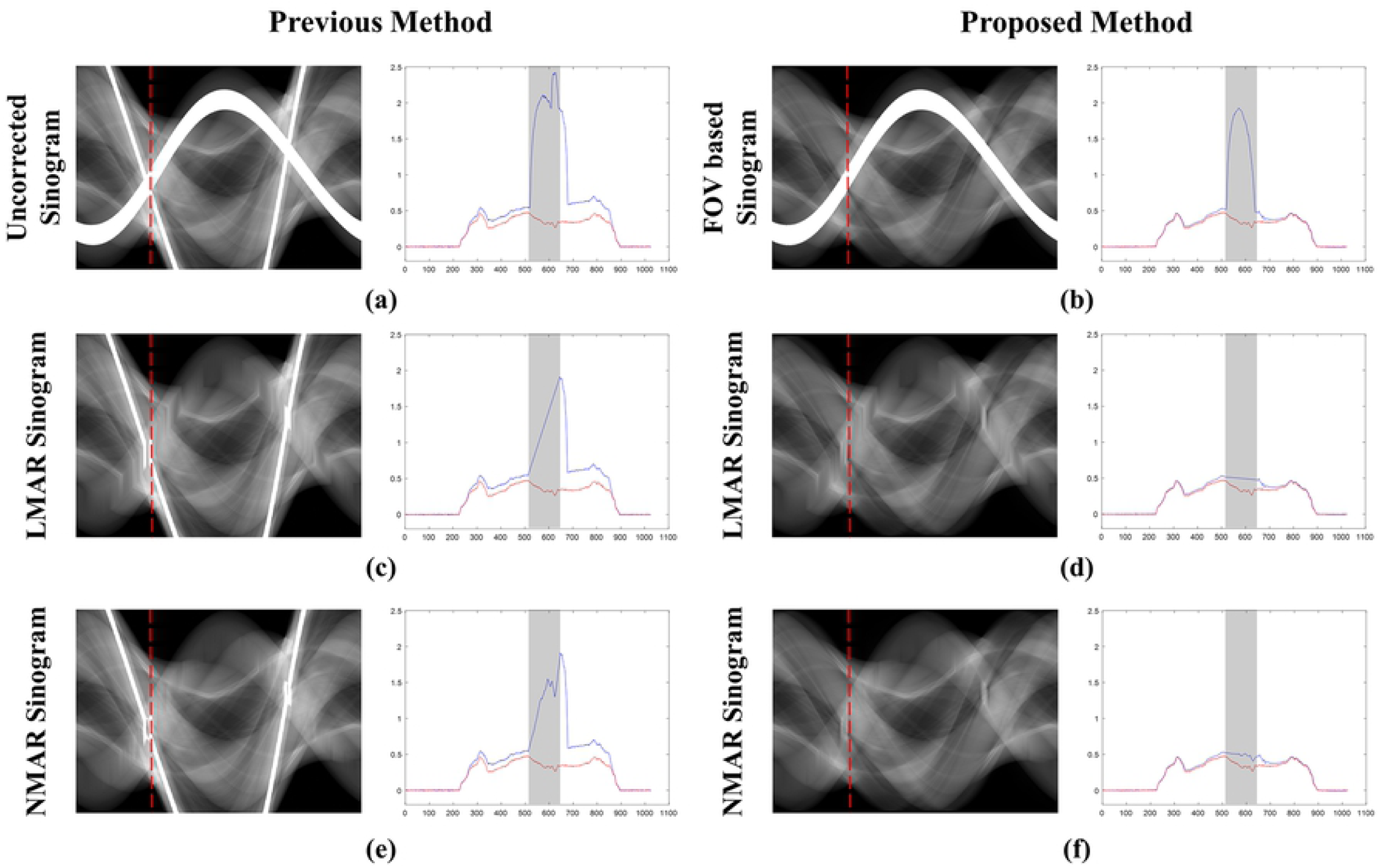
Sinogram and central profile (189th view) of XCAT - shoulder results. (a): Uncorrected sinogram, (b): FOV-based sinogram, (c),(e): sinogram from previous LMAR and NMAR method, (d),(f): sinogram from the proposed LMAR and NMAR method. In the sinogram images, the red dotted lines represent the central profile at the 189th view and the gray shades in the central profiles represent the metal trace region. In the plots, the blue lines represent each sinogram value at the 189th view and the red lines represent the reference sinogram value at the 189th view.

Table 2 summarizes NMSE and SSIM for the LMAR and NMAR results, which are compared with the ROI area of the reference image for each head, shoulder, and hip case. NMSE and SSIM show better performance of the proposed method compared to the previous method. In addition, the NMAR shows better performance compared to LMAR in the proposed method.

**Table 2.**
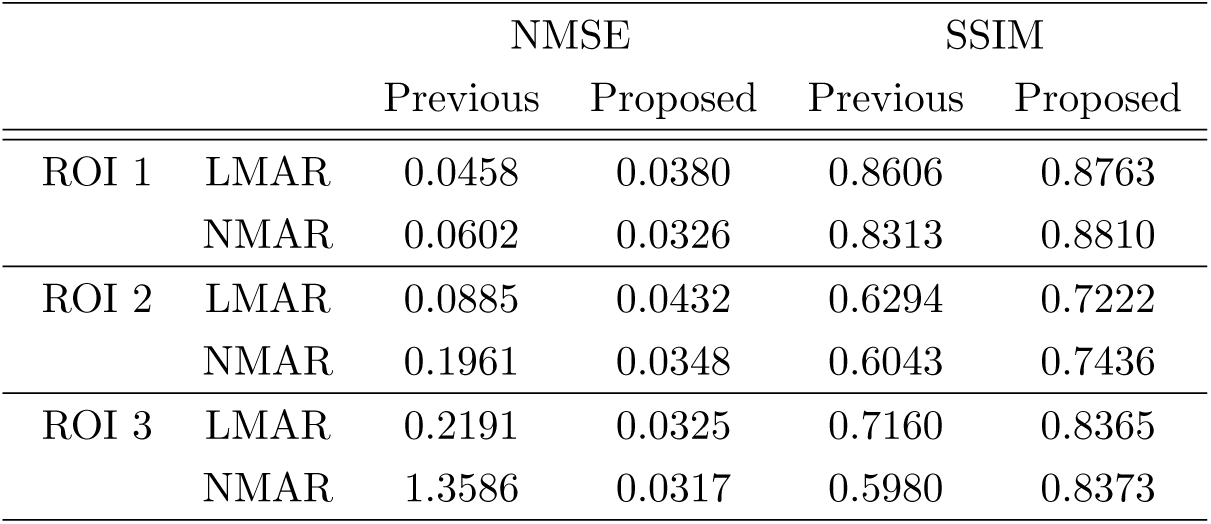
NMSE & SSIM results with XCAT image.

|  |  | NMSE |  | SSIM |  |
| --- | --- | --- | --- | --- | --- |
|  |  | Previous | Proposed | Previous | Proposed |
| ROI 1 | LMAR | 0.0458 | 0.0380 | 0.8606 | 0.8763 |
|  | NMAR | 0.0602 | 0.0326 | 0.8313 | 0.8810 |
| ROI 2 | LMAR | 0.0885 | 0.0432 | 0.6294 | 0.7222 |
|  | NMAR | 0.1961 | 0.0348 | 0.6043 | 0.7436 |
| ROI 3 | LMAR | 0.2191 | 0.0325 | 0.7160 | 0.8365 |
|  | NMAR | 1.3586 | 0.0317 | 0.5980 | 0.8373 |

### Clinical Data Results

Fig. 8 represents the result images for previous and proposed LMAR and NMAR methods for the clinical data of the abdomen, shoulder, hip, and head. The leftmost column is the reference image for each case. For image comparison, the area of the metal is marked in yellow. The red box shows the ROI area for quantitative comparison of image quality. The next two images show the LMAR result, which is the result obtained with LMAR using the previous method on the left and using the proposed method on the right. The last two images are the NMAR results, which are the results obtained from using previous method on the left and the proposed method on the right. The results show that all MAR results reduce the metal artifacts when compared to the uncorrected metal artifact images. However, as indicated by the red arrow, the residual artifacts remain with the previous method, which are effectively removed with the proposed method.

**Fig 8.**
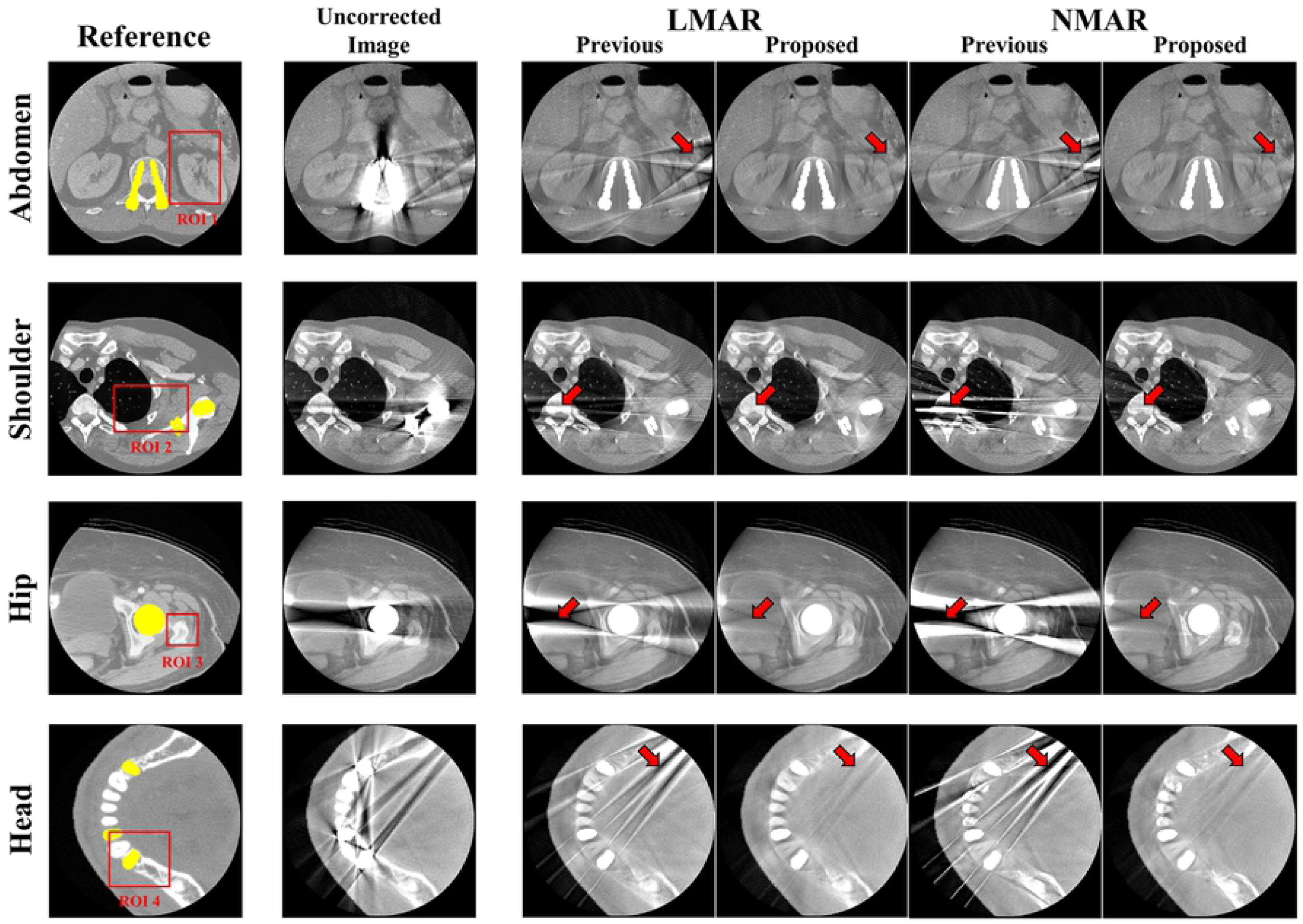
Result with clinical data. Each row corresponds to different body sections. Each column represents reference images, uncorrected metal artifact images, and previous and proposed LMAR and NMAR result images. Window width/window level = 2000HU/0HU.

Table 3 summarizes the NMSE and SSIM for the clinical data. NMSE and SSIM are shown for each abdomen, shoulder, and head case, indicating that the proposed method performs better than the previous method in all cases.

**Table 3.** NMSE & SSIM results with clinical image.

|  |  | NMSE |  | SSIM |  |
| --- | --- | --- | --- | --- | --- |
|  |  | Previous | Proposed | Previous | Proposed |
| ROI 1 | LMAR | 0.0815 | 0.0339 | 0.8413 | 0.9090 |
|  | NMAR | 0.1128 | 0.0286 | 0.8304 | 0.9162 |
| ROI 2 | LMAR | 0.1904 | 0.1632 | 0.7279 | 0.7975 |
|  | NMAR | 0.3824 | 0.1504 | 0.6335 | 0.7986 |
| ROI 3 | LMAR | 0.0597 | 0.0350 | 0.8800 | 0.9298 |
|  | NMAR | 0.1517 | 0.0298 | 0.7423 | 0.9407 |
| ROI 4 | LMAR | 0.0479 | 0.0442 | 0.9040 | 0.9154 |
|  | NMAR | 0.0459 | 0.0339 | 0.8981 | 0.9155 |

### Experimental Results

Fig. 9 shows the original disk phantom, where the number of metals inside the FOV is fixed as three and the number of metals outside the FOV increases from one to three. The MAR results are shown in Fig. 10. In all cases, the MAR results are better than the uncorrected original images. However, as indicated by the red arrows in the MAR image obtained with the previous method, dark streak artifacts still remain. As the number of metals outside the FOV increases, the distortion of the image becomes severe. On the other hand, as indicated by the red arrows in the MAR image obtained with the proposed method, the metal artifacts are effectively reduced. Previous MAR method can reduce the metal artifacts inside the FOV(i.e, streak artifacts between inside metal objects), but residual streak artifacts still remain because the metal artifacts produced by metals outside the FOV are difficult to remove. However, with the proposed method, the FOV-based sinogram mitigates the effects from the external metal trajectory, which reduces the residual metal artifacts more effectively. It is also observed that the NMAR is more effective to recover the boundary of the cylinder phantom than LMAR, as indicated by the green arrows in Fig. 10. Table 4 summarizes the NMSE and SSIM for the experimental results. Quantitative assessments also confirms our observation.

**Table 4.** NMSE & SSIM results with experimental image.

|  |  | NMSE |  | SSIM |  |
| --- | --- | --- | --- | --- | --- |
|  |  | Previous | Proposed | Previous | Proposed |
| 1 Metal | LMAR | 0.2031 | 0.1767 | 0.8142 | 0.8338 |
|  | NMAR | 0.1604 | 0.1304 | 0.8184 | 0.8442 |
| 2 Metals | LMAR | 0.2447 | 0.1997 | 0.7388 | 0.7754 |
|  | NMAR | 0.2099 | 0.1653 | 0.7397 | 0.7819 |
| 3 Metals | LMAR | 0.2702 | 0.2062 | 0.7048 | 0.7554 |
|  | NMAR | 0.2357 | 0.1703 | 0.7089 | 0.7612 |

**Fig 9.**
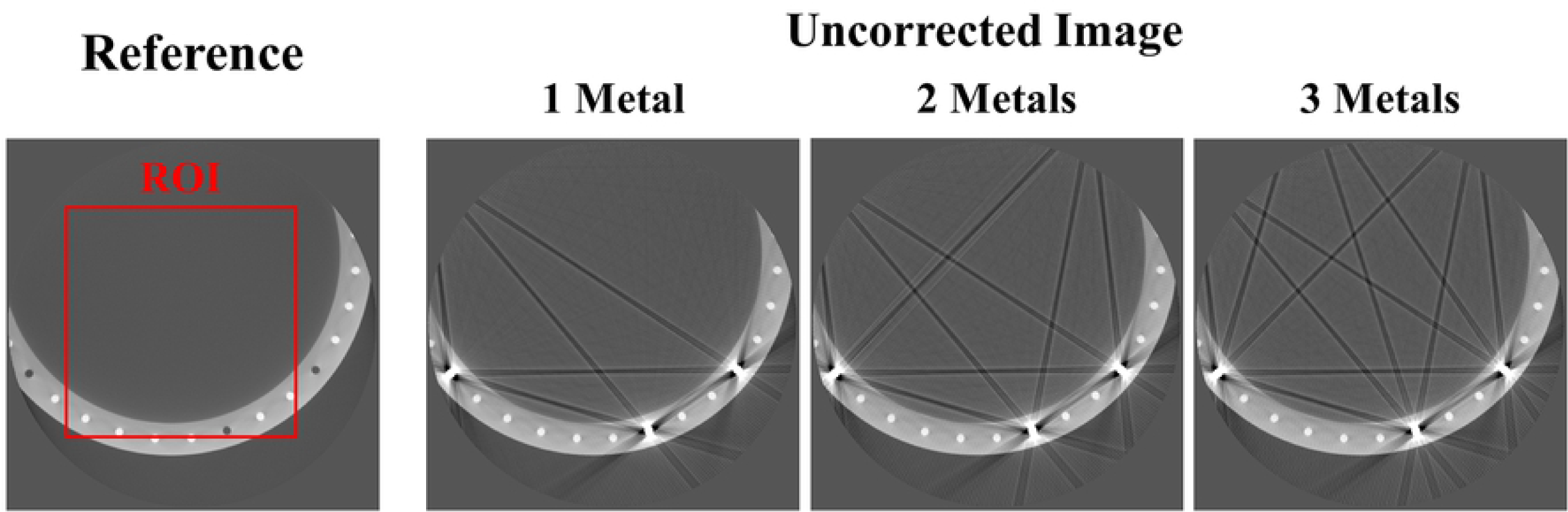
Reference image and uncorrected images from experimental data of the disk phantom. Window width/window level = 3000HU/-500HU. The red box represents the ROI area where the NMSE and SSIM are measured.

**Fig 10.**
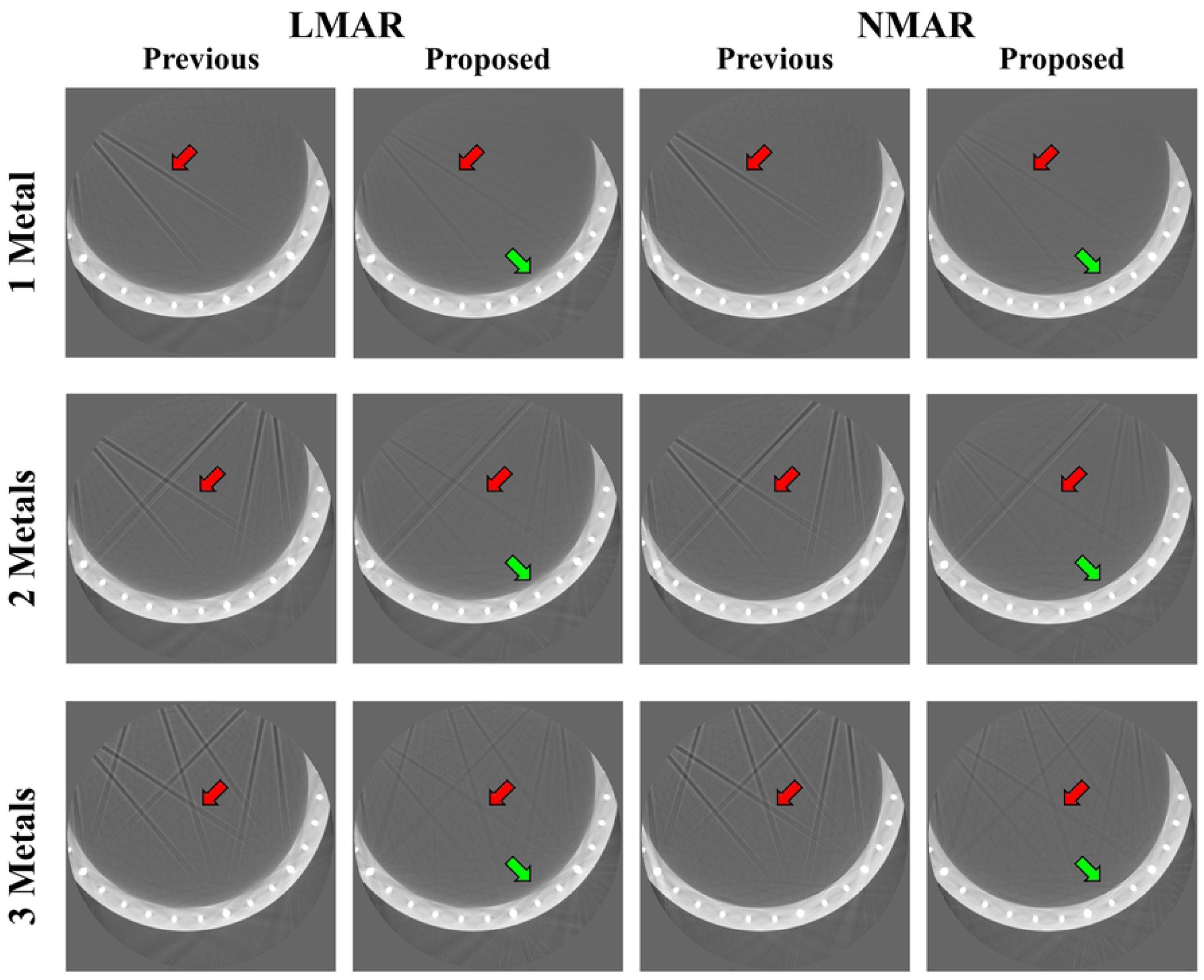
Result of disk phantom experimental data. Each row corresponds to different numbers of metals outside the FOV. Each column represents the LMAR and NMAR result images with the previous and proposed methods. Window width/window level = 2500HU/-750HU.

## Discussion and Conclusion

In sinogram based inpainting MAR methods, using precise values between metal trace regions is critical to estimate the missing data within the metal trace region. However, when metal objects reside outside the FOV, the neighborhood values of the interpolated metal traces are corrupted by these metals, which introduces more interpolation errors, resulting in residual artifacts after MAR. Furthermore, for small FOV imaging, when using a sinogram inpainting method with a prior sinogram like NMAR, the level of the originally measured sinogram is not matched with the forward-projection of the small FOV image, which can introduce additional artifacts after conducting MAR. Using the proposed method, synthesized projection data for a small FOV image are acquired and treated as the originally measured sinogram, which minimizes the effect of the metals residing outside the FOV during the inpainting procedure.

In this paper, only LMAR and NMAR were used to represent the results. Since, the proposed method uses a FOV-based sinogram instead of the original sinogram, the proposed method can be applied for other sinogram inpainting methods such as the metal deletion technique [21] and total variation-based MAR [22, 23]. In addition, the proposed method proceeds through forward-projection of the first reconstructed image, rather than using the original sinogram, and thus the raw data of the CT are not needed for the whole process. Therefore, the proposed method can be used to reduce the metal artifacts in CT images if the information of the data acquisition geometry is provided.

In conclusion, we have proposed a FOV-based MAR method that takes into consideration the presence of data truncation. Instead of using the uncorrected sinogram for MAR, the proposed method proceeds through forward-projection of the uncorrected metal artifact image. The results confirmed that LMAR and NMAR with the proposed method can reduce artifacts generated by metal from both inside and outside the FOV, which could not be removed effectively with the previous method.

## Acknowledgments

This research was supported by the Bio and Medical Technology Development Program of the National Research Foundation (NRF) funded by the Ministry of Science and ICT (NRF-2018M3A9H6081483 and 2019R1A2C2084936).

